# A 3D adrenocortical carcinoma tumor platform for preclinical modeling of drug response and matrix metalloproteinase activity

**DOI:** 10.1101/2023.01.24.525287

**Authors:** Priya H. Dedhia, Hemamylammal Sivakumar, Marco A. Rodriguez, Kylie G. Nairon, Joshua M. Zent, Xuguang Zheng, Katie Jones, Liudmila Popova, Jennifer L. Leight, Aleksander Skardal

## Abstract

Adrenocortical carcinoma (ACC) has a poor prognosis, and no new drugs have been identified in decades. The absence of drug development can partly be attributed to a lack of preclinical models. Both animal models and 2D cell cultures of ACC fail to accurately mimic the disease, as animal physiology is inherently different than humans, and 2D cultures fail to represent the crucial 3D architecture. Organoids and other small 3D *in vitro* models of tissues or tumors can model certain complexities of human *in vivo* biology; however, this technology has largely yet to be applied to ACC. In this study, we describe the generation of 3D tumor constructs from an established ACC cell line, NCI-H295R. NCI-H295R cells were encapsulated to generate 3D ACC constructs. Tumor constructs were assessed for biomarker expression, viability, proliferation, and cortisol production. In addition, matrix metalloproteinase (MMP) functionality was assessed directly using fluorogenic MMP-sensitive biosensors and through infusion of NCI-H295R cells into a metastasis-on-a-chip microfluidic device platform. ACC tumor constructs showed expression of biomarkers associated with ACC, including SF-1, Melan A, and inhibin α. Treatment of ACC tumor constructs with chemotherapeutics demonstrated decreased drug sensitivity compared to 2D cell culture. Since most tumor cells migrate through tissue using MMPs to break down extracellular matrix, we validated the utility of ACC tumor constructs by integrating fluorogenic MMP-sensitive peptide biosensors within the tumor constructs. Lastly, in our metastasis-on-a-chip device, NCI-H295R cells successfully engrafted in a downstream lung cell line-based construct, but invasion distance into the lung construct was decreased by MMP inhibition. These studies, which would not be possible using 2D cell cultures, demonstrated that NCI-H295R cells secreted active MMPs that are used for invasion in 3D. This work represents the first evidence of a 3D tumor constructs platform for ACC that can be deployed for future mechanistic studies as well as development of new targets for intervention and therapies.

**Significance:** The paucity of preclinical research models has contributed to lack of progress in treatment of adrenocortical carcinoma (ACC). Three-dimensional modeling of ACC may provide novel insights for preclinical studies and advance ACC research.

## Introduction

Adrenocortical carcinoma (ACC) is a rare endocrine malignancy that originates within the adrenal cortex.^1^ Surgical resection is the only curative option; however, most patients present with unresectable disease or have recurrence after surgery. Treatments, such as mitotane, etoposide, doxorubicin, and cisplatin, (EDP) and radiation are used to manage advanced disease, but in most cases metastatic disease is fatal within one year despite these treatments.^1-3^ Therefore, there is a critical need to develop new therapies in order to improve outcomes in patients with ACC.

Development of effective therapies relies on availability and adequacy of preclinical models. Two-dimensional (2D) cell lines have been used extensively to model human diseases. The NCI-H295R cell line was generated to model ACC;^4,5^ however, 2D cell lines do not accurately recapitulate heterogeneity found within most ACC tumors. Furthermore, 2D culture conditions fail to mimic *in vivo* topography, diffusion kinetics, availability of adhesion peptides, and cell-cell interactions, which can cause significant genomic and phenotypic shifts.^6,7^ Indeed, therapies developed based on 2D NCI-H295R preclinical studies were predominantly found to have little effect in clinical trials.^8^

Three-dimensional (3D) cell cultures, such as organoids and hydrogel-supported tumor constructs, which incorporate aspects of tumor architecture and microenvironment.^9-11^ demonstrate cellular heterogeneity, spatial organization, cell-cell interactions, and cell-matrix interactions similar to the original tumor. To date, organoids have effectively modeled key developmental and tumorigenic pathways of gut, kidney, liver, lung, retina, brain and other organs.^12-16^ In addition, organoids often recapitulate drug sensitivities observed in patients, with greater accuracy than 2D cultures.^17-19^ Consequently, organoids have become valuable tools in preclinical modeling.^20-22^ Although fetal adrenal organoids have been developed,^23^ to our knowledge, 3D models of ACC have not been described.

To address this need, we employed a hyaluronic acid and gelatin extracellular matrix (ECM) hydrogel approach to generating 3D ACC tumor constructs. This approach has been customized by our team to match mechanical properties of different tissue types,^6,16,19,24-29^ automate tumor construct biofabrication,^25,30,31^ and generate patient-derived tumor organoids (PTOs) from numerous tumor types, including lung, colorectal, melanoma, appendiceal, mesothelioma, sarcoma, and glioma.^17,30,32-36^

Previous studies have established that ACC tumors exhibit dysregulation of the IGF2 and WNT/β-catenin.^37^ Specifically, IGF2, a ligand in the IGF2 signaling pathway, is a growth factor that promotes proliferation and survival and is highly expressed in up to 90% of ACCs.^38,39^ The Wnt pathway is important in adrenal development and tumorigenesis. Binding of Wnt ligand to its receptor leads to cytoplasmic accumulation and nuclear translocation of β-catenin and transcription of target genes. Nuclear β-catenin is seen in 85% of ACC tumors, suggesting activation of the pathway.^40^ In addition, somatic activating mutations in CTNNB1, the gene that encodes β-catenin are common mutations in ACC tumors.^40,41^ The human H295 ACC cell line (generated in 1990)^5^ shows dysregulation of IGF2 and Wnt similar to a subset of ACCs.^40,42^ In addition, steroid factor-1 (SF-1), Melan A, and inhibin-α are expressed in 69-98% of adrenocortical carcinomas and is expressed in the NCI-H295R cell line.^43-47^

In this study, we developed and evaluated a 3D tumor construct model as a complementary model for ACC and compared them to standard 2D cultures. ACC tumor constructs were generated by encapsulating NCI-H295R cells in hydrogel scaffolds. We then characterized ACC constructs and 2D cultures by determining biomarker expression (SF-1, Melan A, inhibin-α, IGF2, and β-catenin), cortisol production, and viability after treatment with standard therapeutic agents, mitotane and the combination of EDP. Finally, we assessed potential utility of 3D ACC constructs over 2D cultures by assessing matrix metalloproteinase (MMP) activity. To demonstrate the potential of 3D constructs in complex modeling, we expanded this MMP activity assessment to a microfluidic model of metastasis, interrogating the roles this enzyme family holds in tissue engraftment and invasion. Our work demonstrated that bioengineered ACC tumor constructs have similar biomarker expression and cortisol production but have reduced drug sensitivity compared to 2D cultures. Furthermore, 3D ACC constructs have measurable MMP activity, and furthermore, quenching this activity impairs their motility in metastatic invasion into target tissue constructs. Together these data suggest that 3D models of ACC can serve as a useful supplement to 2D cultures in ACC preclinical studies.

## Results

### Experimental design

3D platforms that incorporate cell lines have been shown to have biologic and mechanical properties that more closely resemble the primary tumor in a number of cancer types.^6,7,29^ Here, we sought to determine the utility of a 3D platform that incorporates the NCI-H295R cell line as an alternative preclinical model for ACC. We biofabricated hyaluronic acid gelatin hydrogel constructs, which are derived from materials native to the human body, to recapitulate aspects of the 3D microenvironment of *in vivo* tissues. NCI-H295R ACC tumor constructs were generated using this hydrogel construct and deployed to assess proliferation, immunofluorescence, steroid production, drug screening, and MMP expression (**Figure 1)**.

**Figure 1.**
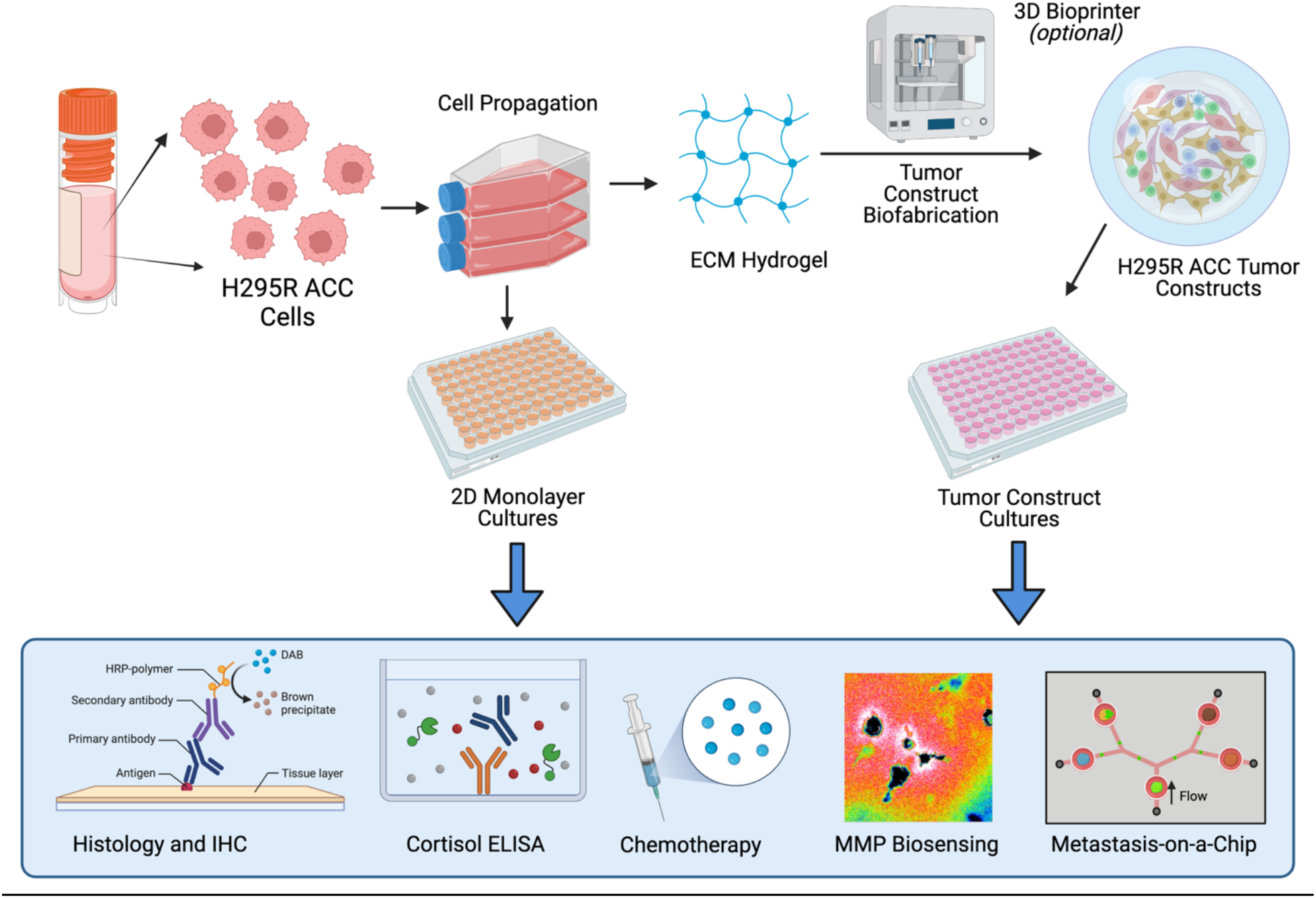
A depiction of the overall workflow of the study. NCI-H295R ACC cells are used either as traditional 2D cell cultures or ECM hydrogel-based 3D tumor constructs. The culture models were then characterized through histology and IF and deployed within chemotherapy screens. The 3D constructs were also functionalized with MMP-sensitive fluorogenic peptide biosensors to evaluate cellular MMP activity within the 3D ECM hydrogel, or implemented within a metastasis-on-a-chip platform.

### 3D ACC constructs express key ACC biomarkers and remain viable in culture

To determine if 3D culture techniques support ACC culture, expression of several ACC protein biomarkers routinely used to diagnose ACC^48,49^ were examined using immunofluorescence. After 7 days of 3D NCI-H295R tumor construct culture, we observed expression of SF-1, Melan-A and, inhibin α are expressed as in 2D culture. As in the majority of ACC, NCI-H295R cells demonstrate IGF2 overexpression and nuclear localization of β-catenin indicating Wnt pathway activation. To determine dysregulation of IGF2 and WNT/β-catenin signaling pathways in NCI-H295R constructs, we examined the expression and localization of IGF2 and β-catenin in 2D cultures and NCI-H295R constructs by immunofluorescence. We observed that these proteins exhibited similar patterns of expression and localization, suggesting that both 2D cultures and 3D NCI-H295R constructs demonstrated dysregulation of IGF2 and WNT/β-catenin (**Figure 2a-j** and **Supplemental Figure 1**).

**Figure 2.**
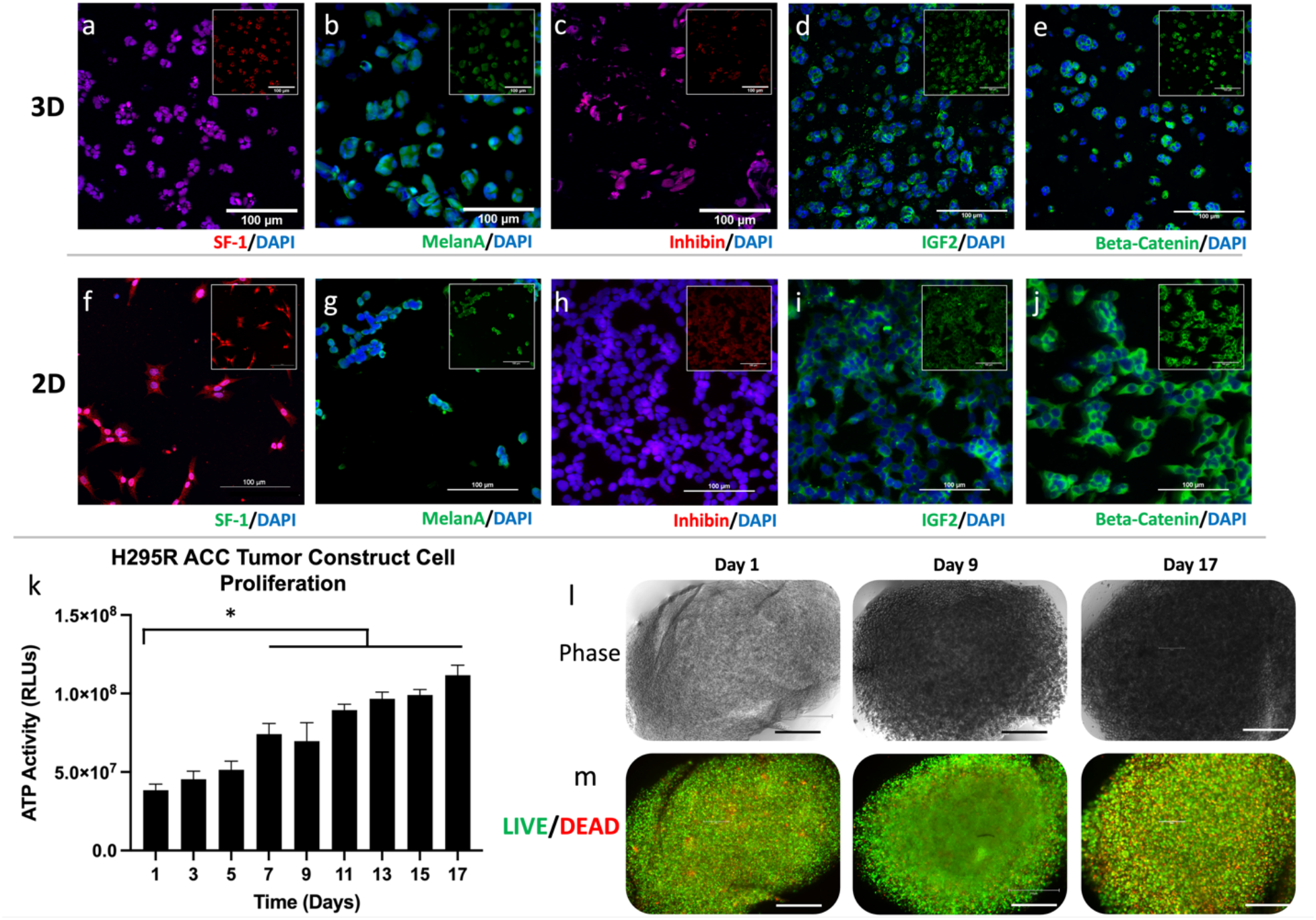
3D ACC constructs express key biomarkers in 3D while proliferating over time *in vitro*. Immunofluorescence staining of key ACC-relevant biomarkers, SF-1, Melan-A, inhibin α, IGF2, and β-cateninin both a-e) 3D construct cultures and f-j) 2D cultures; insets depict stains without DAPI overlay. m) Quantification of ATP activity in 3D ACC constructs over time. Statistical significance: * p < 0.05. k) 3D constructs were maintained for up to 17 days. l) Brightfield imaging demonstrates 3D construct cell density. m) LIVE/DEAD staining shows largely viable cells within 3D constructs over the 17 days of culture. Green – calcein AM-stained viable cells; Red – ethidium homodimer-1-stained dead cell nuclei. Scale bars in panels a-m: 100 μm; n-o): 200 μm.

Subsequently, we assessed ACC construct viability by quantifying ATP activity over a 17-day time course. We observed a consistent increase in ATP activity indicating consistent proliferation of cells within the 3D constructs for the duration of assessment (**Figure 2k**). Brightfield imaging of NCI-H295R constructs demonstrated increased density, also suggesting proliferation of NCI-H295R cells (**Figure 2l**). Lastly, LIVE/DEAD staining – in which viable and dead cells are stained green and red, respectively – and imaging of ACC constructs by fluorescent microscopy indicated that the majority of cells remained viable for 17 days (**Figure 2m**). Together, these data show that NCI-H295R tumor constructs are viable and express ACC biomarkers as in 2D culture.

### 3D ACC constructs increase cortisol secretion in response to forskolin stimulation

Approximately 60% of all ACC tumors produce adrenal hormones. Cortisol, the most common hormone produced, is secreted in 30% - 40% of all functional ACCs.^50^ While cortisol production in adrenal glands is regulated by adrenocorticotropic hormone (ACTH), the NCI-H295R cell line is only modestly responsive to ACTH.^51,52^ However, treating NCI-H295R cell line with forskolin has been shown to induce cortisol release.^53^ In order to evaluate the functional ability of NCI-H295R 3D constructs, we assessed basal and stimulated cortisol production. Both 2D cultures and NCI-H295R 3D constructs demonstrated basal cortisol production, with no significant increase after treatment with ACTH (**Supplementary Figure 2**). However, forskolin stimulation resulted in a significant increase in cortisol production in both 2D cultures and NCI-H295R 3D constructs (**Figure 3**). These data demonstrate that cortisol production is similar between 2D cultures and NCI-H295R 3D constructs. However, we did observe that at the first timepoint (day 0), the 3D constructs did demonstrate significantly higher concentrations of secreted cortisol.We may be able to attribute this to the fact that our ECM hydrogel is a more physiologically accurate substrate for the cells to adhere to through specific integrin-ECM binding compared to the artificial plastic 2D surface on which binding occurs by non-specific hydrophobic bonds. However, over the remaining duration of these experiments, cortisol expression was similar between 2D and 3D groups.

**Figure 3.**
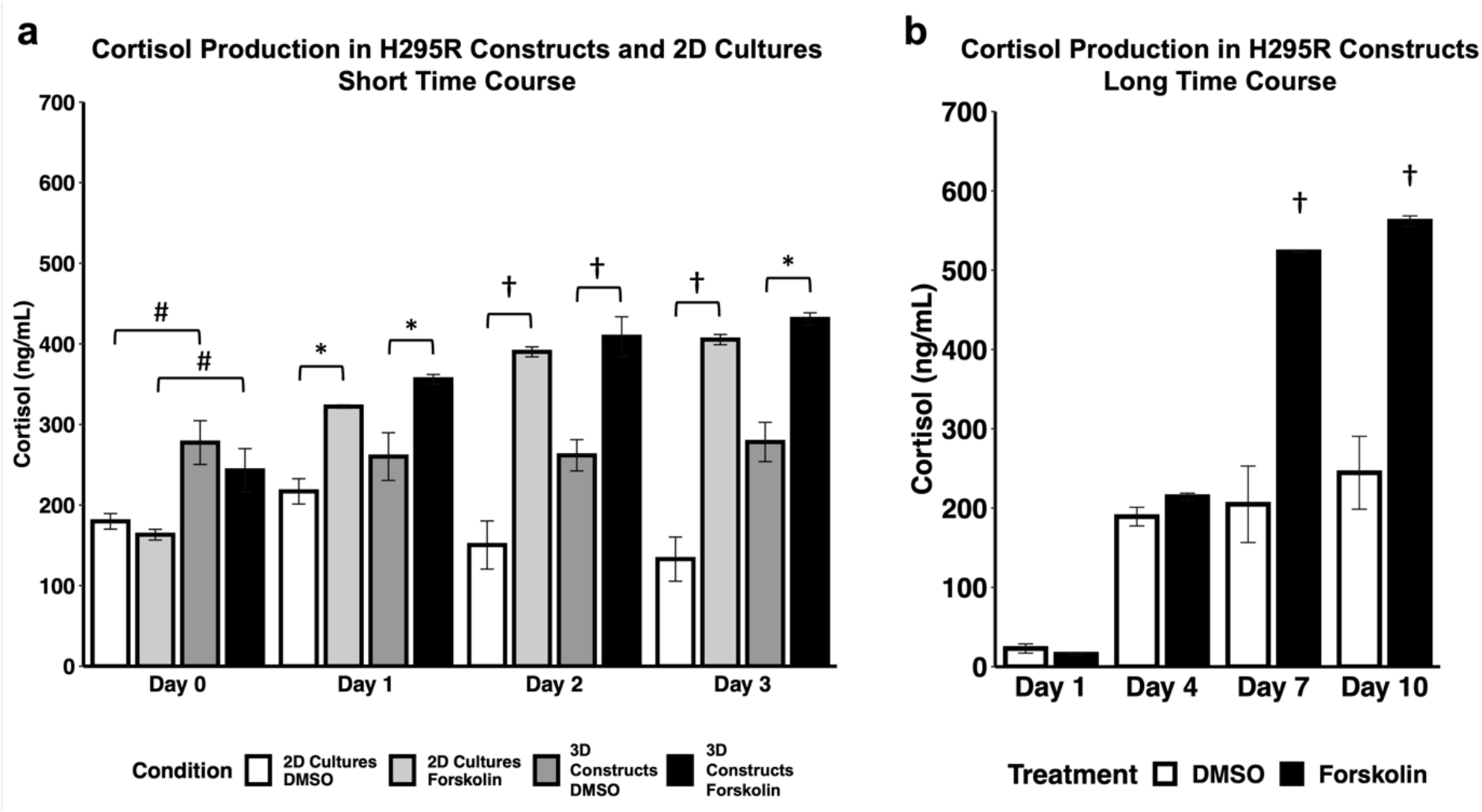
ACC 3D constructs increase secretion of cortisol in response to stimulation. ACC 3D constructs and 2D cell cultures were stimulated with control DMSO or 10 mM forskolin at days 0, 1, 2, and 3 for the short time course (a) and days 1, 4, 7, and 10 for the long time course (b). Cortisol was measured in biological duplicates using enzyme-linked immunoassay. Statistical significance: * p<0.05; † p<0.01 as compared to DMSO control; # p<0.05 between 2D and 3D on Day 0.

### 3D ACC constructs respond to standard ACC chemotherapy regimens

Mitotane is the only drug approved by the United States FDA for treatment of ACC; however, recent studies have shown that only 8% of patients exhibit a complete response when mitotane is prescribed as a single agent.^54^Additionally, mitotane can be combined with other chemotherapeutic drugs such as etoposide, doxorubicin, and cisplatin (EDP) to be administered as part of a treatment regimen. While an EDP plus mitotane regimen is recommended by current ACC treatment guidelines, its efficacy in treating advanced ACC is limited.^55-57^ Prior studies have demonstrated NCI-H295R sensitivity to mitotane.^58,59^ In order to determine if 2D cultures and NCI-H295R 3D constructs would respond differently to treatment with mitotane and the EDP, we treated NCI-H295R 3D constructs with standard chemotherapeutic agents. In 2D cultures, all conditions except the lowest concentration of EDP demonstrated a significant reduction in viability (**Figure 4**). In contrast, only the highest concentration of EDP alone or EDP with mitotane resulted in a significant reduction in viability in NCI-H295R 3D constructs. These data indicate that a 3D platform demonstrates distinct drug-dependent viability compared to 2D cultures.

**Figure 4.**
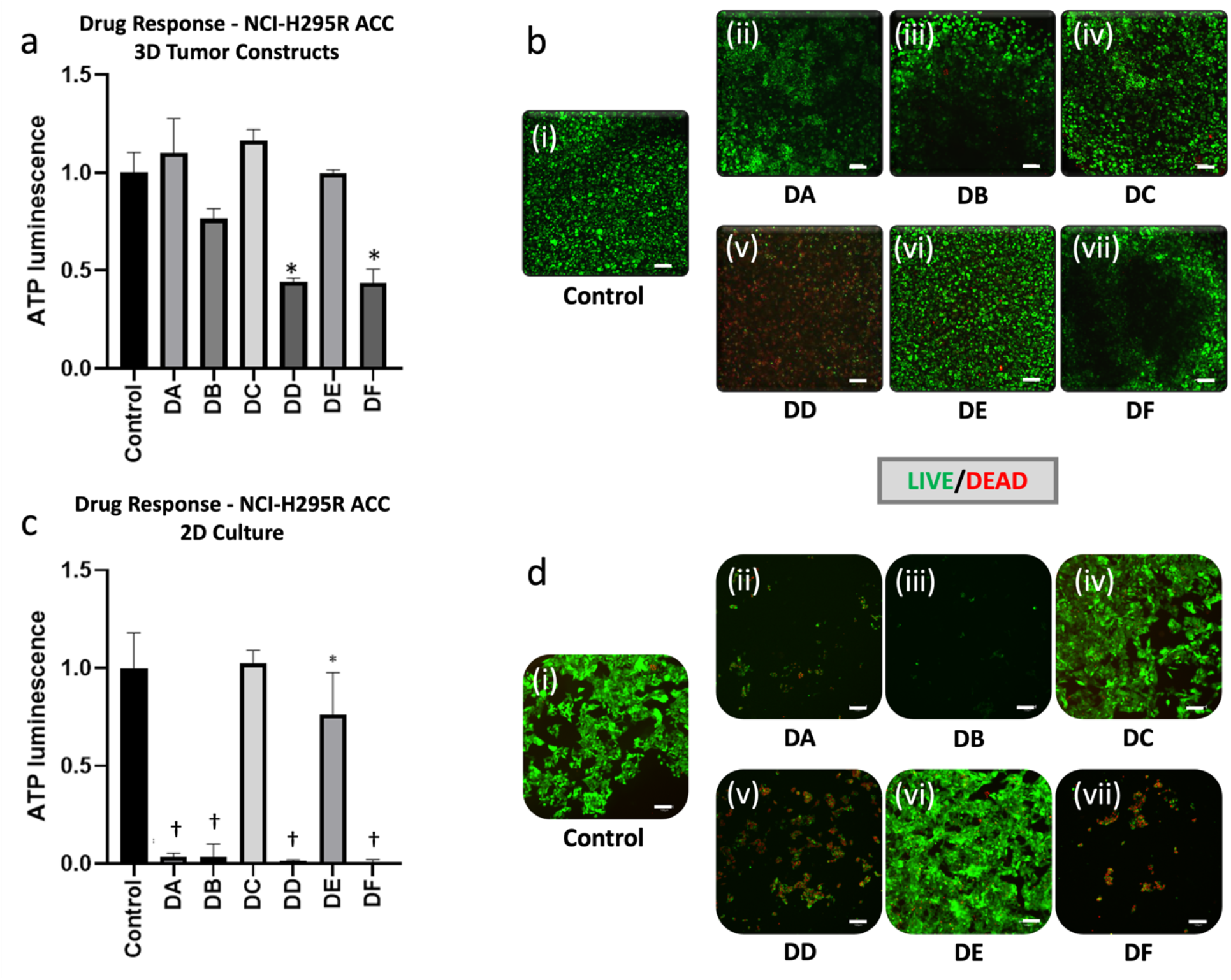
ACC 3D constructs are less responsive than corresponding 2D cell cultures to common ACC chemotherapies. a-b) ACC 3D constructs subjected to drug treatments are evaluated by a) ATP activity quantification and b) LIVE/DEAD staining and fluorescent microscopy. Green – calcein AM-stained viable cells; Red – calcein AM-stained dead cell nuclei. c-d) The same assays, but in 2D cell cultures. Treatment groups are (i) no treatment control, (ii) mitotane at 8 μg/mL (DA), (iii) mitotane at 20 μg/mL (DB), (iv) EDP (etoposide, doxorubicin, and cisplatin) at 0.25 μg/mL, 0.1 μg/mL D, 0.1 μg/mL, respectively (DC), (v) EDP at 25 μg/mL, 10 μg/mL D, 10 μg/mL, respectively (DD), (vi) 8 μg/mL mitotane + EDP at 0.25 μg/mL, 0.1 μg/mL D, 0.1 μg/mL (DE), and (vii) mitotane + EDP at 25 μg/mL, 10 μg/mL D, 10 μg/mL, respectively (DF). Scale bars – 100 μm. Statistical significance: * p<0.05; † p<0.01 as compared to the no treatment control.

### ACC cells actively secrete MMPs within the 3D constructs

Tumor cells migrate using MMPs to cleave ECM components around them thereby creating physical pathways through tissue that are more conducive to movement of the cells. Therefore, we wanted to assay the ability of cells within the 3D tumor constructs to secrete MMPs. An additional benefit of a 3D culture system is the ability to incorporate biosensors within the matrix to measure cellular activity in space and time. Here, we adapted a previously developed MMP biosensor tested extensively using other cancer cells^60-62^ for use in our hydrogel system. The MMP biosensor is comprised of an MMP-degradable peptide sequence based on type I collagen, with a fluorophore and quencher on either side of the MMP cleavage site. Upon cleavage by MMPs, quenching is relieved and fluorescence increases. **Figure 5a** shows the chemistry employed to form 3D tumor constructs with the integrated MMP biosensor. The MMP biosensor was covalently incorporated into the hydrogel system with a light-induced thiol-ene bond between a thiol from a C-terminal cysteine in the biosensor and the methacrylated collagen. After NCI-H295R cells were encapsulated into the biosensor functionalized hydrogels, we evaluated MMP activity through 1) confocal microscopy and 2) by total fluorescence in wells over a range of cell densities. Confocal microscopy in phase shows an NCI-H295R cell (**Figure 5b(i)**), the DiD cell membrane dye fluorescent signal (**Figure 5b(ii)**), and the fluorescence from the QGIW fluorogenic peptide biosensor, indicating MMP expression by the cell (**Figure 5b(iii)**). In addition, **Supplementary Video Files 1** and **2** show 3D projections of the DID cell membrane dye fluorescence and the QGIW fluorogenic peptide biosensor, respectively, on a single NCI-H295R cell. Measurement of global MMP activity using a plate reader demonstrated increased MMP activity with increasing cell density, which was reduced with the addition of the pan-MMP inhibitor GM6001 (**Figure 5c**). These data demonstrate how NCI-H295R 3D constructs may be utilized for studies that are not possible with 2D cell cultures.

**Figure 5.**
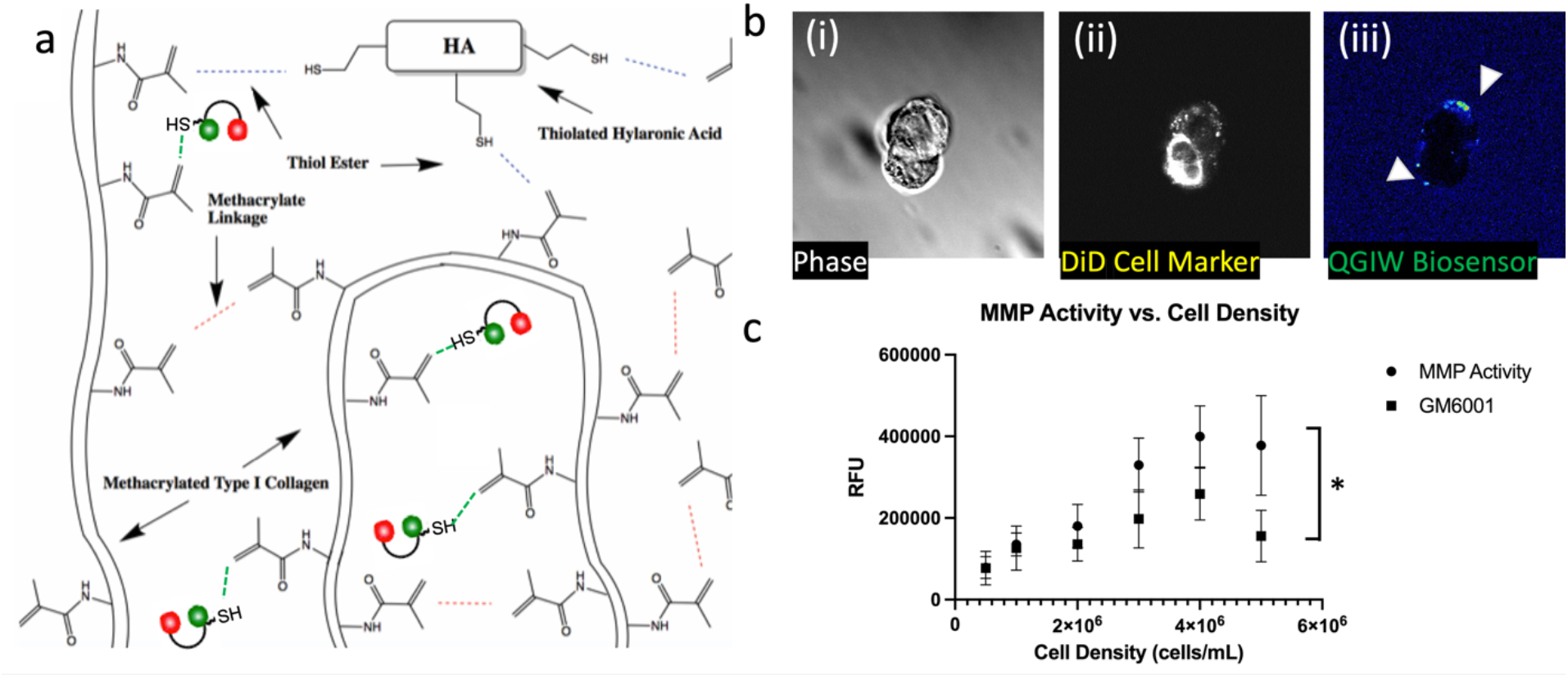
ACC 3D constructs demonstrate MMP activity. a) Light induced thiol-ene bonds between the thiolated HA, thiol group in the MMP biosensor, and methacrylate groups on collagen. b) Biosensor sensing of cell MMP activity in ECM hydrogels using live-cell confocal imaging. (i) Light microscopy of cells within hydrogels. (ii) Fluorescent imaging of the NCI-H295R cell labeled with a DiD cell membrane intercalating dye. (iii) MMP biosensor is visualized by the white signal. c) MMP activity measured in NCI-H295R ACC cell line 3D ECM cultures and with the GM6001 MMP inhibitor. Statistical significance: * p < 0.05.

### MMP activity is required for post-metastasis invasion

To understand the role of MMP activity in ACC invasion, we deployed our MOC platform to mimic metastatic transport and invasion into a metastatic site. As designed, the microfluidic setup (**Figure 6a-b**) mimicked mouse tail vein metastasis modeling using suspended NCI-H295R cells. Tissue constructs of the lung cell line, A549, represented distant sites in our platform.

**Figure 6.**
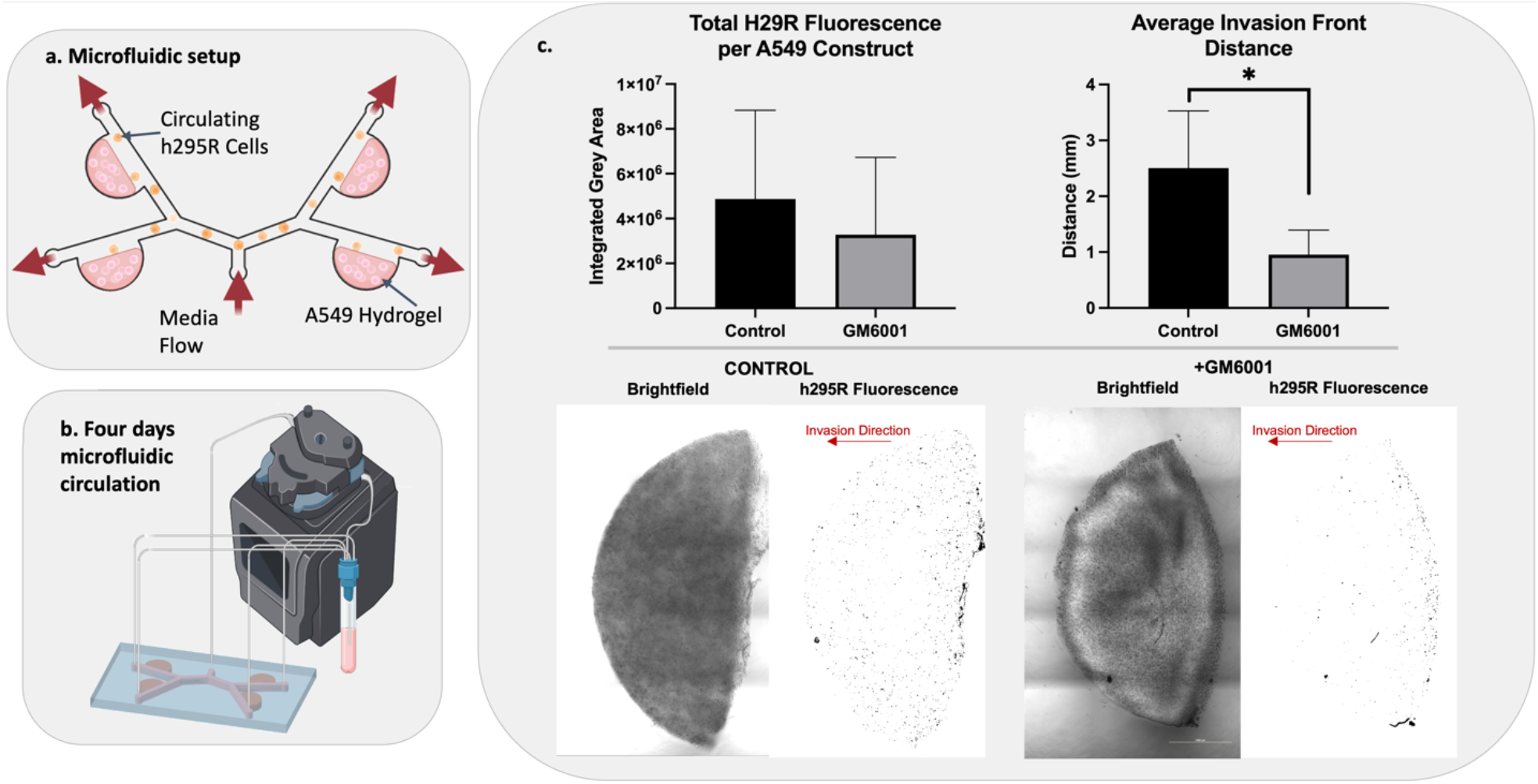
MMP activity is required for invasion. a) Microfluidic devices were designed with four A549 lung epithelial cell-containing hydrogels and a circulating fluorescently-tagged NCI-H295R population. b) ACC cells were circulated through the devices for 4 days either with or without GM6001 MMP inhibitor. c) MMP inhibition did not significantly affect the number of NCI-H295R ACC cells that were able to attach to and invade the A549 lung hydrogels, however the average distance the ACC cells invaded into the A549 lung hydrogels was significantly reduced with MMP inhibition. Representative images demonstrate the disparity in invasion distance between conditions. Statistical significance: * p<0.01

Regardless of MMP inhibition, NCI-H295R cells were able to attach to A549 hydrogels. Total red fluorescence quantification of each A549 hydrogel site showed that MMP inhibition does not have any significant effect on the total number of NCI-H295R cells that leave circulation and begin to invade through the matrix. However, inhibition of MMP activity significantly decreases the distance NCI-H295R cells invade into the hydrogel sites (**Figure 6c**). This implies that initial attachment and extravasation steps in this ACC metastasis model may not be MMP driven, but deeper invasion into the distant site tissues is enhanced by MMP activity. Representative images in Figure 6c show this discrepancy in invasion distance with and without MMP inhibition.

## Discussion

The 5-year survival rate for metastatic ACC is 6%, and current treatments only slow disease progression for 5 months.^1-3^ Thus, there is a critical need to develop targeted therapies for advanced ACC. The startling lack of new targeted therapies is primarily due to the lack of preclinical models that accurately represent ACC, thus significantly limiting evaluation of new drug compounds or new therapeutic strategies. ACC cell lines cultured in 2D fail to recapitulate the 3D complexity of *in vivo* tumors, and available murine models exhibit fundamental differences from human ACC. These shortcomings in current models highlight the real clinical need for improved research models to enhance our understanding of the basic molecular biology of ACC. To address these limitations, here we describe the use of the NCI-H295R cell line in a bioengineered ECM-based 3D tumor construct platform to serve as a more clinically relevant and versatile model system for ACC.

To demonstrate that our ACC tumor constructs accurately represent the disease, multiple characterization assays were performed. We found that the key biomarkers commonly used to verify the identity of ACC cells were present in the 3D ACC constructs (**Figure 2a-e**), similar to 2D cultures (**Figure 2f-j**). These include SF-1, MelanA, inhibin α, IGF2, and β-catenin – which is cytoplasmic and nuclear (**Supplementary Figure 1a-b**). In addition, our data demonstrated continuous cell proliferation and generally high viability. Furthermore, our NCI-H295R constructs demonstrated basal and forkskolin-induced cortisol production similar to 2D culture. Evidence of maintenance of steroidal production in ACC cells *in vitro* is limited, due to the relatively few published *in vitro* ACC studies – all of them being in 2D models.^4,5^ We showed that in both a short 4-day time course and a longer 10-day time course, the addition of forskolin amplified cortisol production in the 3D ACC constructs. Long-term steroid production by ACC cells *in vitro* has been limited to a small number of cell lines. The application of 3D culture of ACC cells may enable longer term steroid production in future patient-derived ACC organoids.

A key reason for developing 3D ACC models is to use them for drug screening assays. ACC 3D tumor constructs can be utilized with cell lines as described herein to test current drugs versus combination treatments, or in the future to test novel drug candidates to aid in drug development efforts. In **Figure 4**, we observe notably different drug responses in our 3D NCI-H295R constructs compared to parallel 2D cultures. ACC constructs were largely resistant to mitotane treatments and showed some response to combinatorial treatment with EDP. On the other hand, 2D cultures were very responsive to mitotane-only treatments, with EDP showing less of an influence in therapeutic efficacy. This discrepancy between 2D and 3D models of ACC is a significant concern in the utility of using 2D models of H295 cell lines alone for clinically relevant research. Mitotane, while a commonly used drug for ACC, has limited efficacy. Here, we have shown that by transitioning from a 2D culture to bioengineered 3D model, we capture mitotane’s variable responsiveness. We should note here that we do not believe the reduced cell death in 3D is due to any limitations on diffusion and penetration of the drug compounds. All of the drugs used in this experiment have molecular weights below 600 g/mol, and thus have very small hydrodynamic radii resulting in increased Brownian motion and efficient diffusion kinetics. In addition, our hydrogels have enhanced diffusion kinetics, as we have demonstrated through drug studies using normal tissue and tumor 3D constructs. In our studies, many treatments are highly effective and can be more effective than in 2D cell cultures, despite needing to diffuse through the hydrogel, even if they have higher molecular weights.^17,18,34,36,63,64^

Metastatic ACC is responsible for a majority of patient deaths. Here we have presented the first in vitro model of ACC metastasis. Our ACC MOC is slightly simplified compared to our previous MOC models in that the microfluidic device houses a target tissue construct, after which tumor cells are infused into the recirculating device, thus mimicking a tail vein injection model. While many mechanistic pathways contribute to metastasis, MMP activity is a key regulator of migration and invasion of tumor cells during metastasis.^65^ MMP expression is elevated in most human tumor types, including breast, colon, pancreas, and melanoma, and increased expression correlates with higher tumor grade and worse prognosis.^66^ MMPs are key regulators of the extensive remodeling of ECM proteins in the tumor microenvironment. This remodeling allows tumor cells to escape and colonize distant sites and promotes blood vessel infiltration. To evaluate the implications of MMPs in ACC, which has been significantly understudied,^67^ we deployed a fluorogenic MMP-sensitive peptide biosensor that was integrated into the 3D ACC construct ECM hydrogels. By using this biosensor-functionalized 3D tumor construct platform, we could directly view MMP activity through fluorescent microscopy. Imaging of the peptide biosensor showed positive expression of MMP-based cleavage of the biosensor (**Figure 5b**). We also used bulk biosensor fluorescence readings in 96-well plates on a plate reader, which showed that the biosensor fluorescence-based MMP activity was significantly decreased by treatment of the ACC constructs with the pan-MMP inhibitor GM6001. The impact of this inhibition was reflected in a significant decrease in GM6001-treated NCI-H295R cells’ ability to invade deep into model lung constructs in a preliminary metastasis-on-a-chip platform. However, we should note that there was not a significant difference in the number of cells that arrived at the lung constructs, regardless of GM6001 treatment or not. As such, we interpreted differences in invasion kinetics as a readout representing one aspect of the metastatic process. In future work, we plan to utilize MOC devices in which the ACC cells are housed in a primary site tumor construct, and need to intravasate into the circulation before they can travel to the lung constructs and engraft. This would be a more complex, yet easy to operate, metastasis model. Nonetheless, these data suggest that MMPs may be involved in ACC progression and metastatic colonization, however, further studies are needed to investigate this.

Both the 3D ACC tumor constructs alone, and the ACC MOC platform have potential for additional translational research. Perhaps most obvious is the deployment of the 3D ACC tumor constructs in drug screening applications. There are only several drugs that are typically used in the clinic to treat ACC patients, and they generally are not effective.^1-3^ Existing drugs used in other cancers, as well as experimental drug compounds, can be tested in the ACC tumor construct platform for tumor cell killing efficacy by targeting alternative pathways. However, we acknowledge that some of our observations in this study are limited by the use of one cell line for these experiments. This is an unfortunate problem facing ACC researchers. Few human cell lines exist. Future work in the ACC research field should be targeted towards determining the viability and clinical relevance of primary patient-derived ACC organoids – work that our team has moved into, that is an extension of our work with patient-derived tumor organoids in other cancers.^17,34,36,64,68-70^ Such patient-derived tumor models can better retain the heterogeneity observed in tumors in vivo, and thus will serve as more appropriate systems in which to screen novel therapeutics. Our MOC platform to date has primarily been used to model metastasis kinetics, largely without any therapeutic intervention.^6,71^ However, as we demonstrate in this study, a treatment intervention – in this case MMP inhibition – reduced the extent of metastatic invasion in the 3D lung constructs. We should note that pan-MMP inhibitors did enter clinical trials, and were shown to be ineffective as clinical therapeutics.^72^ Nonetheless, our study shows that the MOC can certainly be deployed in drug intervention studies, with mitigation of metastasis kinetics being a key output metric in addition to potential tumor cell killing. Future work is needed to determine if targeted MMP inhibition or MMP inhibition in combination with other targeted therapies may play a role in ACC treatment.

ACC represents a disease for which few experimental models exist. Here, we describe the first cell line-based ECM hydrogel 3D tumor construct model that recapitulates distinguishing biomarkers of ACC. Our 3D model better represents the clinical scenario of mitotane resistance, while showing the potential for combinatorial treatments. Our platform also incorporates the capability to measure MMP activity, a primary driver of metastasis. In the future, we hope to use ACC clinical biospecimens to create analogous patient-derived tumor organoids that will be deployed for precision oncology studies.

## Materials and Methods

### Cell culture

NCI-H295R cells (ATCC) at passages 5-8 were initially expanded and cultured in 2D plastic according to ATCC recommendations using DMEM:F12 medium containing 0.00625 mg/ml insulin, 0.00625 mg/ml transferrin, 6.25 ng/ml selenium, 1.25 mg/ml bovine serum albumin, 0.00535 mg/ml linoleic acid, and 2.5% Nu-Serum I (media supplements available from Corning). At 80% confluence, cells were harvested with 0.25% (W/V) Trypsin-0.53 mM EDTA solution (Thermo Fisher) and resuspended in media for 2D culture or 3D hydrogels.

### ACC tumor construct biofabrication

Components of HyStem-HP kit (Advanced Biomatrix) – thiolated and heparinized hyaluronic acid (HA), thiolated gelatin, and the crosslinker polyethylene glycol diacrylate (PEGDA – were dissolved in degassed water containing 0.01% w/v photoinitiator (2-Hydroxy-4′-(2-hydroxyethoxy)-2-methylpropiophenone [Sigma]) at 1%, 1%, and 2% w/v, respectively, and added to pelleted NCI-H295R cells at a 2:2:1 volume ratio with a final cell concentration to 10,000 cells/μl. Hydrogel precursor-cell suspensions of 10 μl were pipetted into 48-well plates that had been coated with thin layers of polydimethylsiloxane to create hydrophobic surfaces, after which each deposited volume was crosslinked via 365 nm UV LED light (BlueWave RediCure 365, Dymax, Torrington, CT) exposure for 2 seconds, and complete media was added. The 10 μL tumor constructs, which remain stable in size for the duration of the experiment, were cultured in 300 μL of media per well, with media aspirated and replenished daily to maintain nutrient supply.

### Histological and immunofluorescence characterization

Immunofluorescence (IF) was performed on sectioned paraffin-embedded tumor constructs or fixed cells cultured for 3 to 7 days on X-tra® slides to analyze biomarker presence of SF-1 (ab240394, Abcam), Melan A (ab210546, Abcam), inhibin-α (ab47720, Abcam), IGF-2 (ab226989, Abcam), and β-catenin (ab ab223075, Abcam). Nuclei were stained with DAPI (P36962, ThermoFisher). The slides were imaged with Nikon A1R confocal microscope at 40X magnification at room temperature.

### Cell proliferation in 3D tumor constructs

Tumor construct cell viability and proliferation were measured over a 17-day period to quantify and visualize NCI-H295R growth in the hydrogel system. ATP and LIVE/DEAD analyses were carried out every other day up to day 17. Cell viability via ATP quantification was carried out using the Promega® CellTiter-Glo® 3D - Superior Cell Viability Assay according to the manufacturer’s instructions. LIVE/DEAD-staining and fluorescent microscopy imaging was carried out using the Viability/Cytotoxicity kit for mammalian cells (Invitrogen) according to the manufacturer’s instructions.

### Steroid production

Cortisol production in response to treatment with forskolin (Sigma-Aldrich) or adrenocorticotropic hormone (ACTH) (Thermo Fisher) was measured in NCI-H295R constructs and 2D cultures. Tumor constructs were prepared as described above and cultured for 4 days. On the fourth day, tumor construct media was collected and replaced (n=3) (Day 0, Figure 3), and forskolin or ACTH was added at concentrations of 10 μM or 100 nM. Media for all groups was collected and replaced (with forskolin, ACTH, and DMSO) on days 1, 2, and 3 post compound addition for the short time course, and days 1, 4, 7, and 10 for the long-time course (all groups n=3). For 2D culture, 100,000 cells were plated into culture treated 48 well plates. This cell number was selected to have the same number of cells per well as the 3D tumor constructs. Cortisol was quantified by human Cortisol ELISA kit (cat. no. ab108665; Abcam) according to manufacturer’s instructions.

### Drug screening in 3D tumor constructs and 2D cultures

NCI-H295R tumor constructs and 2D cultures (50,000 cells) were exposed to mitotane (S1732, Selleckchem, Houston, TX), etoposide (S1225, Selleckchem), doxorubicin (S1208, Selleckchem), and cisplatin (S1126, Selleckchem). The tumor constructs or 2D cell cultures were exposed to clinically relevant drug conditions of mitotane (8 μg/mL or 20 μg/mL), a combination of etoposide, doxorubicin, and cisplatin (EDP) at concentrations of 0.25 μg/mL, 0.1 μg/mL D, 0.1 μg/mL, respectively, or at concentrations of 25 μg/mL, 10 μg/mL D, 10 μg/mL, respectively. In two drug conditions, mitotane at 8 μg/mL was combined with EDP at 0.25 μg/mL, 0.1 μg/mL D, 0.1μg/mL or mitotane at 20μg/mL and EDP at 25μg/mL, 10 μg/mL D, 10 μg/mL respectively. After 7 days for 3D tumor constructs or 3 days for 2D cultures, media was replaced with fresh media containing the aforementioned drug concentrations. Viability was assessed at 72 hours using ATP quantification and LIVE/DEAD staining as described above. For the 2D drug study, NCI-H295R cells were plated at 50,000 cells per well in 48-well plates for 72 hours in drug-free media and at 70% confluence subjected to the drug screen for 2 days as described above to evaluate the effects 2D tissue culture conditions on drug response. ATP quantification and LIVE/DEAD staining were employed as described above.

### Matrix metalloproteinase biosensor assays

The fluorescent MMP-degradable peptide biosensor [GGPQG↓IWGQK_AdOO_C] (where ↓ indicates the MMP cleavage site) was synthesized using Fmoc solid-phase peptide synthesis (Liberty Blue™ Peptide Synthesizer; CEM, Matthews, NC) with a Rink Amide MBHA resin (EMD Millipore, Burlington, MA) as previously described.^61^

For cell encapsulation, hydrogel components were added to the cell solution for a final concentration of 2 mg/ml thiolated hyaluronic acid (THA), 250 μM of MMP biosensor, and 20 mg/ml lithium phenyl-2,4,6-trimethylbenzoylphosphinate (LAP). After addition of collagen methacrylate (6 mg/ml, CMA) and CMA neutralization buffer, 20 μl of cell-gel solution per well was pipetted into ethanol sterilized 96-well black U-bottom plate (BrandTech #781607). Hydrogels were photopolymerized with 4 mW/cm^2^ of 365 nm UV light for 180 seconds, and media was added. For MMP inhibition, GM6001 (abcam ab120845) was diluted to a final concentration of 20 μM. After 16 hours of culture at 37°C for 16 hours, biosensor fluorescence was measured at an excitation of 485 nm and an emission of 528 nm using a microplate reader (Synergy H1) with a 5 x 5 area scan for a total of 25 points measured per well.

### Confocal microscopy of MMP biosensor

Cells were prepared as for MMP biosensor then were stained with 5 μg/ml of DiD cell tracker (Invitrogen D7757). Cells were encapsulated at a density of 2.5x10^6^ cells/ml and biosensor concentration of 50 μM. For imaging studies, hydrogels were fixed to Ibidi glass bottom μ-dishes (Ibidi #81158). To facilitate hydrogel attachment through thiol-ene covalent bonding, μ-dish glass was thiolated by treatment with 3-mercaptopropyl-triethoxysilane (Sigma #175617). Hydrogels were photopolymerized with 4 mW/cm^2^ of 365 nm UV light for 90 seconds and serum-free media was added. Cells were imaged at 48 hours with a Nikon A1R inverted confocal microscope in resonance-scanning mode using a 60x oil objective (1.4 N.A, WD 0.13 mm) with 64x line averaging at a nominal resolution of 0.21 μm/pixel. The biosensor channel was detected using a 487 nm laser with a 535/50 filter and DiD and differential interference contract channels were obtained with a 638 nm laser.

### Metastasis-on-a-Chip

#### Microfluidic Fabrication

To mimic tail vein injection style metastasis models, microfluidic devices were designed with a single inlet port branching to four discrete channels running parallel to four hydrogel chambers. Molds for the upper portion of the devices were laser cut from polymethylmethacrylate (PMMA) sheets, around which polydimethylsiloxane (PDMS) prepolymer (Sylgard 184, Dow Corning, Midland, MI) was poured. After degassing to remove air pockets and curing overnight at 60°C, the PDMS layers were detached from their molds and irreversibly bonded to glass slides activated with air plasma at 1.5 torr for 90 seconds.

#### Cell Embedding and Metastasis Studies

A549 lung epithelial cells were used as a model of lung tissue for potential metastatic invasion. A549 cells were suspended at 10,000 cells/μL in a 3:1 solution of methacrylated collagen I and thiolated hyaluronic acid. Using 30 insulin needles, the precursor solution was injected into all four microfluidic hydrogel chambers and crosslinked in situ for 2 seconds using a 365 nm UV light. NCI-H295R cells were fluorescently tagged using MemGlow 560 (Cytoskeleton, Inc., Denver, CO) following manufacturer instructions. Following this, the cells were suspended in media at 1.67 x 105 cells/mL. Cell suspensions were transferred into media reservoirs, which were then connected to inlet and outlet ports in the microfluidic via a microperistaltic pump (Elemental Scientific, Inc., Omaha, NE) and flow was maintained throughout the device at 10 μL/min for 4 days. For MMP inhibition studies, pan-MMP inhibitor GM6001 (Sigma Aldrich, St. Louis, MO) was added to media reservoirs at 20 μM. At the end of the circulation period, fluorescence and brightfield imaging were performed using a Nikon A1R Live Cell Imaging Confocal Microscope (Nikon Corp., Tokyo, Japan) to quantify NCI-H295R invasion. Z-stacks of 100 μm were taken for each hydrogel construct, from which 2D maximum projections images were formed. Fluorescent images were thresholded using ImageJ software to select for invaded NCI-H295R cells, and brightfield images were used to create a mask defining the hydrogel perimeter.

## Statistical Analysis

Statistical procedures and graph plotting were conducted in GraphPad Prism 6.01 (GraphPad Software) and R version 4.1. All tumor construct studies used sample size of n = 3-5. Statistical analysis was performed using Student’s t-test or ANOVA test. All of the results are presented as mean with error bars representing standard deviation (SD).

## Author Contributions

PD, JL, and AS conceived the study concept and design, obtained funding, and supervised the studies. HS performed drug treatment studies and histological/immunofluorescence staining assays. MR performed tumor construct culture proliferation assessment and collection of tumor construct conditioned media for steroid quantification. XZ performed ELISAs. LP performed statistical analyses. JZ performed matrix metalloproteinase biosensor studies. KN and KJ performed invasion assays. All authors contributed to manuscript and approved the submitted version.

## Data Availability

The datasets used and/or analysed during the current study available from the corresponding author on reasonable request.

