## Supplemental materials for "A 3D adrenocortical carcinoma tumor platform for preclinical modeling of drug response and matrix metalloproteinase activity"

**Supplementary Figures**

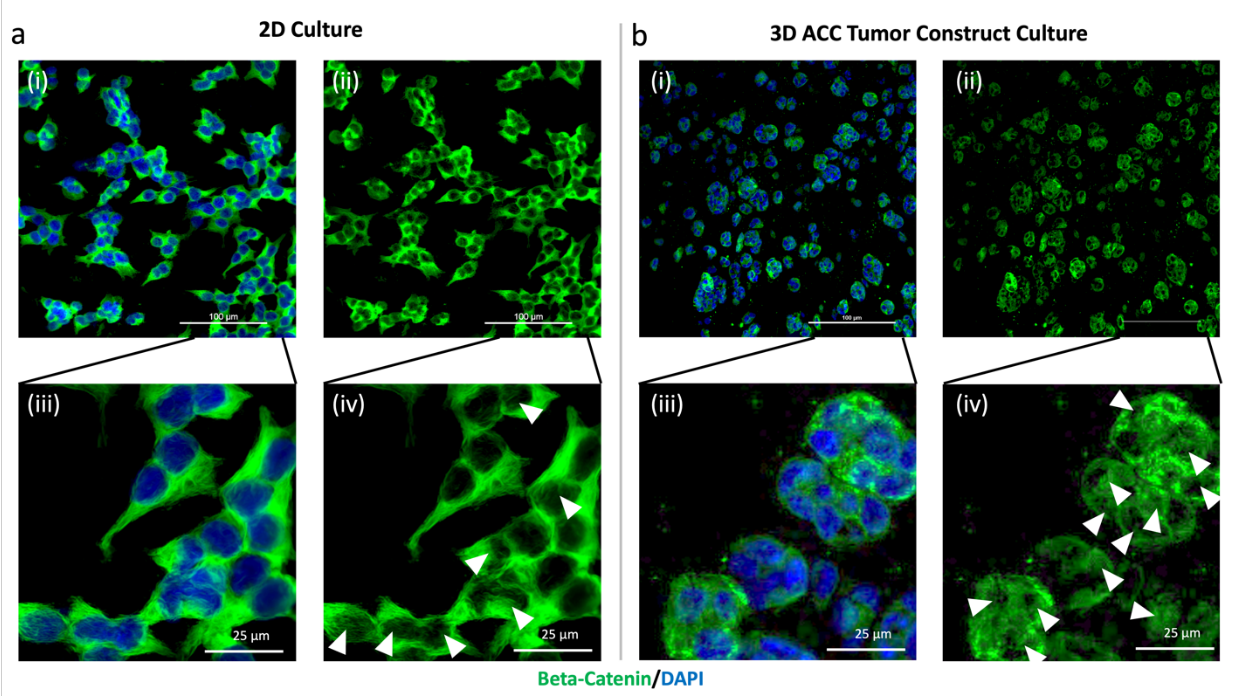


**Supplementary Figure 1. β-catenin expression is found in the cytoplasm and nuclei indicating activation of the Wnt pathway in 2D and 3D cultures.** Fluorescent imaging of NCI-H295R ACC cells cultured a) on 2D tissue culture plastic or b) in 3D tumor construct cultures. For both a) and b) panels, (i) shows the β-catenin expression (green) with stained nuclei (DAPI-blue) and (ii) β-catenin expression only (green). Analogous higher resolution images (iii) show β-catenin expression (green) with stained nuclei (DAPI - blue) and (iv) β-catenin expression only (green). Both cytoplasmic and nuclear β-catenin is observed in both 3D and 2D conditions. Arrows – positive nuclear β-catenin. Scale bars: 100 µm for (i) and (ii) panels; 25 µm for (iii) and (iv) panels.

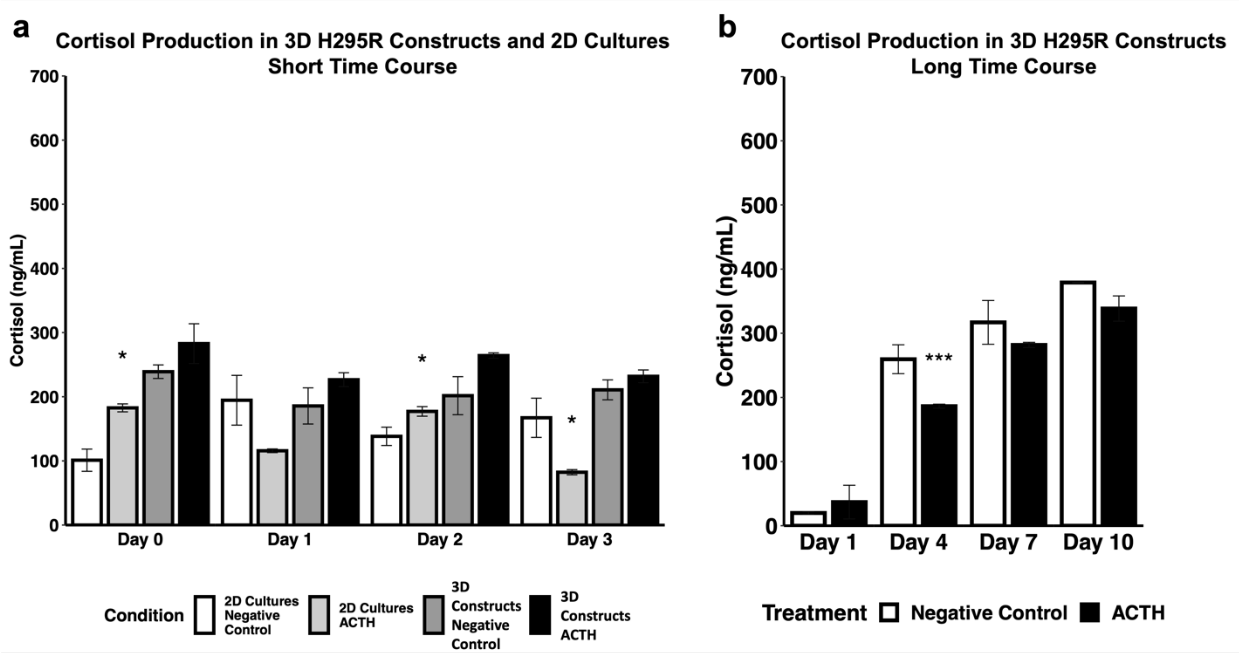


**Supplementary Figure 2. 3D ACC tumor constructs do not increase cortisol secretion in response to ACTH.** 3D ACC tumor constructs and 2D cell cultures were stimulated with media control or 10 nM ACTH at days 0, 1, 2, and 3 for the short time course and days 1, 4, 7, and 10 for the long time course. Cortisol was measured in biological duplicates using enzyme-linked immunoassay. Statistical significance: * $p<0.05$; † $p<0.01$.
